# Tissue-resident CD8^+^ T cells drive age-associated chronic lung sequelae following viral pneumonia

**DOI:** 10.1101/2020.04.13.040196

**Authors:** Nick P. Goplen, Yue Wu, Youngmin Son, Chaofan Li, Zheng Wang, In Su Cheon, Li Jiang, Bibo Zhu, Katayoun Ayasoufi, Eduardo N. Chini, Aaron J. Johnson, Robert Vassallo, Andrew H. Limper, Nu Zhang, Jie Sun

## Abstract

Lower respiratory viral infections, such as influenza virus and severe acute respiratory syndrome coronavirus 2 (SARS-CoV2) infections, often cause severe viral pneumonia in aged individuals. Here, we report that influenza viral pneumonia leads to chronic non-resolving lung pathology and exaggerated accumulation of CD8^+^ tissue-resident memory T cells (T_RM_) in the respiratory tract of aged hosts. T_RM_ accumulation relies on elevated TGF-β present in aged tissues. Further, we show that T_RM_ isolated from aged lungs lack a subpopulation characterized by expression of molecules involved in TCR signaling and effector function. Consequently, T_RM_ cells from aged lungs were insufficient to provide heterologous protective immunity. Strikingly, the depletion of CD8^+^ T_RM_ cells dampens persistent chronic lung inflammation and ameliorates tissue fibrosis in aged, but not young, animals. Collectively, our data demonstrate that age-associated T_RM_ cell malfunction supports chronic lung inflammatory and fibrotic sequelae following viral pneumonia in aged hosts.

## Introduction

Aged individuals (65 and older) will comprise 20 percent of the population in developed countries in the coming decades. It is well established that the peripheral CD8^+^ T cell compartment that contributes to cellular immunity shrinks in number, diversity, and quality as we age [1-3]. As a consequence, memory T cells are enriched and T Cell Receptor (TCR) repertoires are narrowed during homeostasis [4]. Consequently, the magnitude and the quality of primary effector CD8^+^ T cell response often is compromised in aged hosts following pathogen challenge [5]. Likewise, the development of CD8^+^ memory T cell responses in the circulation and/or in the lymphoid organs is often impaired during aging [6, 7]. This age-related CD8^+^ T cell attrition in memory T cell differentiation and/or function has been linked to impaired host responses to pathogens following primary infection or vaccination in aged hosts [8].

Tissue-resident memory CD8^+^ T cells (T_RM_) with potent effector potential are mainly located in peripheral tissue at portals of pathogen entry, whereas circulating central and effector memory T cells survey lymphoid organs and/or peripheral tissues [9, 10]. It has been shown that CD8^+^ T_RM_ cells can provide nearly sterilizing protection if present in sufficient numbers [11]. Therefore, great enthusiasm has recently centered on enhancing local CD8^+^ T_RM_ responses as a promising vaccination strategy to control mucosal infections including influenza virus infection [12]. Much progress has been made recently regarding the cellular and molecular mechanisms regulating T_RM_ cell development and maintenance [13, 14]. However, relatively less is known regarding the effects of aging in the development and/or function of T_RM_ responses in the mucosal tissues following primary infection. Of note, analysis of spatial and temporal maps of T cell compartmentalization revealed that aged lungs harbor increased levels of CD8^+^ CD69^+^ T_RM_-like cells compared to young lungs [1]. However, it is unclear whether this is caused by a gradual build-up of memory T cells following frequent experiences of prior infections in the lungs, or due to enhanced generation of CD8^+^ T_RM_ cells following a single infection in the elderly. Therefore, the development and function of CD8^+^ T_RM_ cells in aged hosts following primary infection remains to be examined.

Lower respiratory tract viral infections represent a major public health challenge and economic burden worldwide. In a matter of months, emerging respiratory viral diseases can alter social norms, stagnate economies, and overwhelm healthcare infrastructures across the globe, as demonstrated by the ongoing severe acute respiratory syndrome coronavirus 2 (SARS-CoV-2) pandemic. Among respiratory viral pathogens, influenza virus can infect 5–10% of the population [15], resulting in up to 35.6 million illnesses and 56,000 deaths annually in the U.S. alone since 2010 [16]. For reasons that are still not fully defined, many respiratory viruses including SARS-CoV-2 and influenza virus cause disproportionately severe morbidity in the elderly population [17, 18]. For instance, the vast majority of the mortality associated with both influenza and SARS-CoV-2 infections, occur in people 65 years and older. It has been shown that influenza disease severity is elevated, prolonged, and linked with increased comorbidities and death in the elderly population [19]. In addition to the acute mortality caused by respiratory viral infections, evidence has suggested that survivors of primary viral pneumonia may display persistent impairment of lung function due to lung fibrosis [20]. Furthermore, it is predicted that a large proportion of elderly Corona Virus Disease 2019 (COVID-19) survivors will be prone to persistent impairment of lung function and pulmonary fibrosis as was observed in survivors of SARS and Middle East Respiratory Syndrome (MERS) [21]. At this time, there are no preventative means or therapeutic interventions available to mitigate and/or reverse lung fibrosis development following viral pneumonia. Currently, the cellular and molecular mechanisms regulating the development of chronic lung fibrotic sequelae following primary viral pneumonia are still largely unknown. This major knowledge gap likely represents a future bottleneck for development of successful therapeutics for patients with chronic lung fibrosis induced by SARS-CoV-2 infection.

Here, we report that influenza infection in aged mice leads to non-resolving inflammation and persistent chronic lung pathology. Transcriptional and flow cytometric analyses revealed that lungs from aged mice exhibited enhanced accumulation of CD8^+^ CD69^+^ memory T cells. Parabiosis experiments showed that those age-associated lung CD8^+^ CD69^+^ T cells were tissue resident. Further, the excessive accumulation of lung CD8^+^ T_RM_ cells depends on the aging-associated increase in environmental TGF-β. Single cell (sc) RNA-seq analysis demonstrated that T_RM_ against a major influenza protective epitope exhibited diminished effector function in response to TCR signaling. As a result, CD8^+^ T_RM_ failed to provide protective immunity against a secondary heterologous virus infection in aged mice. Strikingly, we further showed that the depletion of CD8^+^ T_RM_ cells resulted in decreased tissue inflammation and lowered lung collagen deposition. Thus, we have discovered an age-associated paradox in CD8^+^ T_RM_ cell responses. CD8^+^ T_RM_ cell accumulation in aged hosts was not coupled to enhanced protective immunity, but rather, to chronic lung pathology and fibrotic sequelae following primary viral pneumonia.

## Results

### Non-resolving tissue inflammation and pathology following influenza virus infection in aged hosts

To investigate the short and long-term effects of aging on influenza disease severity, young (2 month old) and aged (21-22 month old) C57BL/6 mice were infected with influenza A virus (IAV; A/PR8/34 strain (PR8), Figure 1A). While 100% of young mice survived from intranasal infection with PR8, ∼50% of aged mice succumbed to the infection by day 12 (Figure 1B), which are consistent with previous reports that aged mice exhibited enhanced host mortality following primary influenza infection [22]. Aged mice that survived exhibited prolonged weight recovery, although their weight completely recovered by 20 days post infection (d.p.i.) (Figure 1C). Interestingly, these differences in acute morbidity and mortality were seen despite similar viral replication at 9 d.p.i. and complete viral clearance in the respiratory tract in both groups by 15 d.p.i. (Figure 1D) similar to previous reports [23].

**Figure 1.**
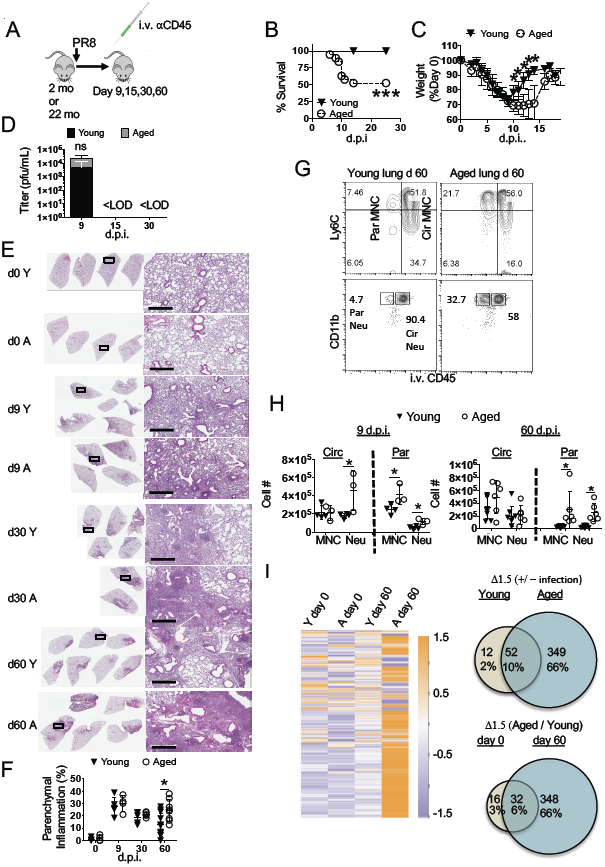
Aged lungs display non-resolving pulmonary immunopathology following influenza infection. Young (2 month) or aged (22 month) C57BL/6 mice were infected or not with PR8 virus. **F-H**, anti-CD45 was injected intravenously (i.v.) to label circulating white blood cells prior to sacrifice. **A.** Schematics of experimental procedure. **B.** Percent of host survival following influenza infection in young or aged mice. **C.** Percent of pre-infection body weight following influenza infection in young or aged mice. **D.** Airway viral titers were determined through plaque forming unit (pfu) assay of BAL fluid at 9, 15, and 30 d.p.i. **E.** Lung histopathology by H & E staining at 0, 9, 30, and 60 d.p.i. **F.** Percent of left lung lobe parenchyma infiltrated by white blood cells was quantitated by Image J analysis. **G, H.** Lung cells were stained for monocytes (MNC: CD64^-^ Siglec-F^-^ CD11c^-^ MHCII^-^ CD11b^hi^) and neutrophils (Neu: Ly6G^+^) which were enumerated in the lung and divided into circulating (Cir) or parenchymal (Par) following intravenous labeling of immune cells at 9 and 60 d.p.i. **G.** Representative plots. **H.** Cell numbers of circulating and parenchymal monocytes and neutrophils at 9 and 60 d.p.i. **I.** 530 transcripts were examined at the endogenous mRNA levels by nanostring from young or aged lungs prior to (day 0), or 60 d.p.i. and displayed as a heatmap (purple = relatively low expression, orange = relatively high expression) following log2 transformation of raw expression data above detection threshold. Venn diagrams displaying data in both number of genes and % of total genes examined for the ratios infected versus not by age (top right panel) or aged to young by day post-infection (bottom right panel) from nanostring data. Groups from **B & C** are n=10 young and n=14 aged mice (2 experiments pooled). **D** is 6 BAL samples per group (2 experiments pooled) for days 9 & 15 and 3 each for day 30. **D, E, G**, and **I** are representatives of 2-4 experiments each; **H** is 1-2 independently significant replicates pooled. Scale bars on histology figures are 600 μm. * p<0.05 Student’s two-tailed t-test with unequal variance.

Recent clinical and experimental evidence suggests that severe acute influenza infection may cause persistent lung pathology, remodeling, and pulmonary dysfunction [20, 24]. To determine whether these acute differences in host morbidity result in differential tissue pathological sequela, we examined the kinetics of histopathology in the lungs following acute PR8 infection in young and aged hosts. While common features of pulmonary aging such as enlarged alveoli were observed prior to infection (0 d.p.i.), leukocytic infiltrates were not prominent in either aged or young naive lungs (Figure 1E). At peak of host inflammatory responses and weight-loss (9 d.p.i.) and even several weeks later (30 d.p.i.), lung histopathology and tissue damage appeared comparable between young and aged mice (Figure 1E & F). However, at 60 d.p.i., lungs from aged mice showed a higher density of parenchymal disruption compared to lungs from young mice (Fig. 1E & F). These data suggest that acute influenza virus infection in aged mice results in persistent non-resolving pathological pulmonary responses.

To confirm these lung histopathological data, we examined the presence of inflammatory monocytes and neutrophils, two major inflammatory cells that can cause pulmonary inflammation and tissue damage [22, 25]. To discern the anatomical location of the cell infiltrates, we examined the vascular and parenchymal fractions of the lungs through the injection of intravenous CD45 antibody (Ab) 5 min before sacrifice [26] (Figure 1G). In this setting, CD45 _i.v_ ^-^ cells were within lung tissue, while CD45 _i.v_ ^+^ cells were in lung blood vessels. Neutrophil counts in the circulation exhibited a slight increase in aging relative to young as has been reported at 9 d.p.i. (Figure 1H) [22]. Further, there were more tissue-resident neutrophils and monocytes in the aged group at 9 d.p.i. (Figure 1H). By 60 d.p.i., the differences of neutrophil and monocytes numbers in the circulation were lost compared to 9 d.p.i., yet evidence of parenchymal neutrophil and monocytes persisted in aged lungs compared to young mice (Figure 1H). Together, the data point to an infection-induced age-related non-resolving inflammatory response following primary influenza virus infection.

We next measured the expression of 560 immune-related genes in the lungs before (0 d.p.i.) or at 60 d.p.i. using Nanostring. The immune gene profiles of the lungs from young or aged uninfected mice were quite similar except a few genes-associated with “inflammaging” [27], which were modestly increased in aged lungs (Figure 1I). Further, the expression profile of the immune-related genes in the lungs from infected young mice (60 d.p.i.) was quite similar to those of lungs from uninfected mice, suggesting that lungs from infected young mice largely return to immune homeostasis at 60 d.p.i. However, the expression of most immune response-related genes were upregulated in infected aged lungs relative to those of uninfected young or aged lungs, or infected young lungs (Figure 1I). The mRNA signal ratios of infected versus non-infected lungs in the young or aged groups identified 52 differentially expressed genes (DEGs) shared between young and aged groups; 12 were unique to young and 349 were unique to aged samples (Figure 1I, top right panel). When comparing gene expression ratios of aged to young lungs, before and after infection, there were only 16 unique age-related DEGs prior to infection, whereas 32 were shared prior to and after infection between young and aged groups. In contrast, 348 unique DEGs were observed at day 60 post-infection in aged groups compared to infected young lungs (Figure 1I, bottom right panel). Among those DEGs, influenza infection led to greatly increased expression of myeloid cell-associated genes, cytokine and chemokine genes, and genes involved in lymphocyte responses in aged lungs (Figure S1A-C). Together, these data indicated that the immune landscape of aged and young lungs is grossly similar prior to infection, but aged lungs exhibit greatly exacerbated inflammatory gene expression and failed to return to immune homeostasis months after primary infection.

### Excessive accumulation of CD69^+^ parenchymal memory CD8^+^ T cells in aged hosts

To gain more insight on the lung global transcription profiles of influenza-infected young or aged mice, we performed bulk RNA-seq on lung tissues from infected young or aged mice at 60 d.p.i. There were around 700 genes differentially expressed between the lungs of young and aged mice (Figure 2A). Among those 700 DEGs, aged lungs expressed higher levels of inflammatory cytokines and molecules-associated with adaptive immune responses (Figure S1D). Gene set enrichment analysis (GSEA) found that young lungs were enriched with gene sets associated with tissue repair such as epithelial-mesenchymal transition (EMT) and apical junction (Figure S1E). In contrast, aged lungs were enriched with gene sets associated with inflammatory responses and adaptive immune responses (Figure 2B & C and S1F).

**Figure 2.**
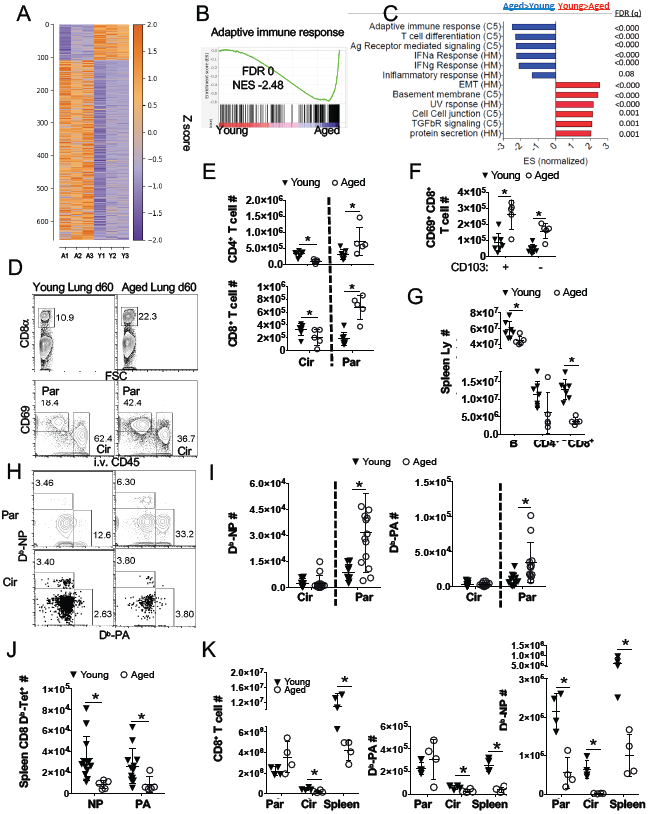
Aged lungs exhibit enhanced local adaptive immune responses despite circulatory deficiencies in the memory phase. Young and aged mice were infected with PR8. **A.** Bulk RNAseq was performed on young or aged lungs at 60 d.p.i., data is displayed as a heatmap (purple = relatively low expression, orange = relatively high expression) of 658 differentially expressed genes (549 up-regulated in aged relative to young, 109 up-regulated in young relative to aged) with p < 0.001. **B.** GSEA for adaptive immune response genes with normalized enrichment score (NES) and false discovery rate (FDR). **C.** Ranked normalized geneset enrichment scores (ES) from the C5 and Hallmark (HM) databases and associated false-discovery rates (FDR (q)). **D.** Representative flow cytometry plot showing lung CD8^+^ T cells separated by CD69 and CD45 i.v. labeling, showing parenchymal (Res) and circulating (Cir) populations as indicated. **E.** Quantification of CD4^+^ (top panel) and CD8^+^ T cells (bottom panel) in lung circulation (Cir) or parenchyma (Par) 60 d.p.i. **F.** CD69^+^ parenchymal CD8^+^ T cells that either express CD103 (CD103^+^) or not (CD103^-^) were enumerated at 60 days post-infection. **G.** T (CD4^+^ & CD8^+^) and B lymphocytes (B220^+^) were quantitated in spleens at 60 d.p.i. (Spleen Ly #). **H.** Representative flow cytometry plots, and **(I.)** number of D^b^-NP (top) or D^b^-PA tetramer positive (bottom) circulating (Circ) or parenchymal (Res) memory CD8^+^ T cells. **J.** Number of D^b^-NP or D^b^-PA tetramer positive CD8^+^ memory T cells in spleens. **K.** Total CD8^+^, CD8^+^ D^b^-NP or CD8^+^ D^b^-PA T cell numbers were quantitated in the lung parenchyma (Par), vasculature (Cir) or spleen (Spleen) at 10 d.p.i. **D-F, H, & K** are representatives of 2-4 experiments. **I & J** are pooled data from 3 independently significant experiments. * p<0.05 Student’s two-tailed t-test with unequal variance.

We were intrigued with the enrichment of adaptive immune responses in the aged lungs as previous reports described that influenza infected aged mice had decreased levels of memory T cell responses in the circulation [28]. Therefore, we performed flow cytometric analysis to measure splenic, lung circulating and parenchymal memory T cell responses following i.v. injection of CD45 Ab as above at 60 d.p.i. (Figure S2A). There were increased CD4^+^, CD8^+^ T and B cell presence in aged lung parenchyma but not within the lung vasculature (circulation) (Figure 2D & E and Figure S2B & C). Lung CD8^+^ CD69^+^ CD103^-^ or CD8^+^ CD69^+^ CD103^+^ T_RM_-like cells were also increased in aged lungs compared to those of young lungs (Figure 2 D, F). Consistent with previous reports, adaptive immune cells, particularly CD8^+^ T cells, were decreased in the spleens of aged mice [29] (Figure 2G).

We next examined influenza-specific CD8^+^ memory T cells in the lungs and spleens of infected young or aged mice through the staining of H2D^b^/nucleoprotein (NP) peptide 366-374 tetramer (D^b^-NP) or polymerase (PA) peptide 224-233 (D^b^-PA) tetramers (Figure 2 H and Figure S2 D). Aged lungs harbored significantly higher numbers of influenza-specific D^b^-NP and D^b^-PA memory T cells, selectively in the lung tissue, compared to young lungs (Figure 2I). In contrast, there was a drastic reduction of influenza-specific CD8^+^ memory T cells in aged spleens (Figure 2J). Thus, we observed an unexpected increase of memory T cells present in aged lungs, despite the attrition of memory T cells in the spleen or circulation. Increased T_RM_ responses in the lungs could be due to increased infiltration of effector T cells in aged mice. We therefore examined effector T cell responses in the lungs and spleens at 10 d.p.i. We found that while D^b^-PA responses appear to be equivalent in the lungs between young and aged mice, there was a deficit in the generation of D^b^-NP specific CD8^+^ T cells in the lungs (Figure 2K). Furthermore, there were significantly lower levels of both D^b^-NP and D^b^-PA effector T cells in the spleens and circulation (Figure 2K) of aged mice compared to controls as previously reported [33]. Therefore, aged mice exhibit enhanced CD8^+^ T_RM_ numbers despite having lower levels of effector T cell responses.

### Influenza-specific lung memory CD8^+^ T cells of aged hosts are tissue resident

To verify whether the CD8^+^ CD69^+^ T cells in the lung that were protected from intravenous labeling where bona fide tissue-resident cells, we infected aged mice with influenza and performed parabiosis surgery to join the circulation of infected aged mice (CD45.2^+^) with a young naive congenic (CD45.1^+^) animal at 5 weeks post infection (Figure 3A). Four weeks after parabiosis, we observed equilibration of CD8^+^ T cells from each parabiont in the blood and spleens (Figure 3B & C and S3A), confirming the successful exchange of circulating immune cells between parabionts. A vast majority of tissue-resident alveolar macrophages belonged to the host in both animals as would be expected from this population that is maintained mainly through self-renewal [34] (Figure S3B). Together, these two observations indicate that shared circulation was achieved and the model is able to distinguish the origin of circulating and residential immune cells in a manner consistent with previously described findings.

**Figure 3.**
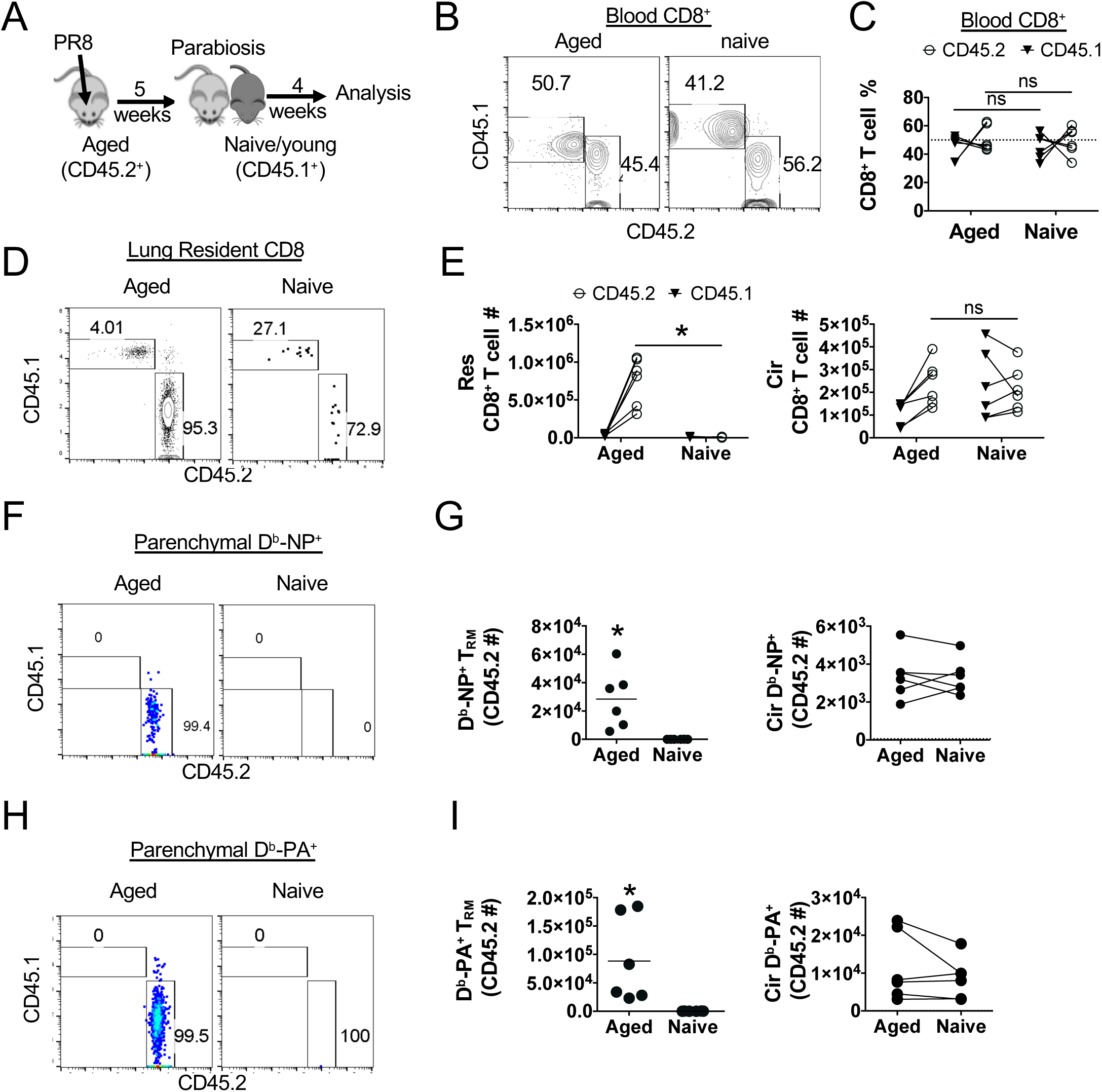
CD8^+^ memory T cells in the lung parenchyma from aged mice are tissue resident. Aged mice (CD45.2) were infected with PR8 and parabiosed with young naive (CD45.1) mice at 5 weeks after infection. 4 weeks later, CD45^+^ white blood cells were intravenously labeled in each host before sacrifice. **A.** Schematics of experimental procedure. **B.** Each animal was bled at time of sacrifice and CD8^+^ T cells were examined by flow cytometery for origin (representative flow plot). **C.** Percent of host or donor CD8^+^ T cells in aged or naive hosts where each line represents all the CD8^+^ T the cells in blood from one mouse. **D.** Representative flow plot of total parenchymal CD8^+^ T cells (CD69^+^CD8^+^ T cells protected from i.v. CD45 labeling) derived from host or donor. **E.** Enumeration of lung parenchymal (left panel) or vasculature (right panel; circ) CD8^+^ T cells in each host by mouse of origin. **F.** Representative flow plot of parenchymal D^b^-NP tetramer^+^ CD8^+^ T cells (CD69^+^CD8^+^ T cells protected from i.v. CD45 labeling) derived from host or donor. **G.** Enumeration of D^b^-NP^+^ cells that are resident (left panel) or circulating (right panel). **H.** Representative flow plot of parenchymal D^b^-PA tetramer^+^ CD8 T cells (CD69^+^CD8^+^ T cells protected from i.v. CD45 labeling) derived from host or donor. **I.** Enumeration of D^b^-PA^+^ cells that are resident (left panel) or circulating (right panel). Experiment was repeated twice with 3 pairs each and data was pooled. * p<0.05 Student’s two-tailed t-test with unequal variance.

The parenchymal compartment of the aged infected hosts had a significantly larger number of CD8^+^ CD69^+^ T cells than the uninfected parabiont, of which ∼95% were from the aged parabiont, despite indistinguishable numbers of CD8^+^ T cells in both the lung vasculature and spleens between the young and aged parabionts (Figure 3D & E). Roughly 99% of total parenchymal CD8^+^ CD69^+^ D^b^-NP^+^ cells in the lungs of two parabionts were found in aged lungs (Figure 3 F, G). In contrast, there were comparable CD45-i.v.^+^ CD8^+^D^b^-NP^+^ cells that were found in the lung circulation between young and aged parabionts (Figure 3G). Likewise, we observed similar patterns on the distribution of the D^b^-PA-specific CD8^+^ T cells in lung parenchyma and circulation (Figure 3H & I & S3E). Notably, there were no major differences in CD8^+^ splenic memory T cells between the two parabionts for the same antigen specificities (Figure S3D & F). Together, these data suggest that aged lung parenchymal CD8^+^ influenza-specific memory T cells are mainly tissue-resident.

### Age-associated T_RM_ cell accumulation is dependent on environmental TGF-β

It remains unclear if the excessive accumulation of T_RM_ cells in aged lungs is due to the aged environment or intrinsic T cell differences in the development and/or repertoire between young and aged mice. To examine these possibilities, we adoptively transferred CD8^+^ Ovalbumin (OVA)-TCR transgenic OT-I T cells of young naive donors to young or aged hosts. We then infected the mice with recombinant influenza virus expressing cognate OVA peptide (SIINFEKL) (PR8-OVA) [35] and evaluated memory T cell responses at 7 weeks after infection (Figure 4A). There were increased numbers of OT-I T_RM_ present in aged lung tissue versus young (Figure 4B), while equivalent numbers of lung circulating or splenic memory OT-I cells were observed between young and aged mice (Figure 4C & D). Together, these data indicated that the aged microenvironment facilitates lung T_RM_ cell accumulation following influenza virus infection even though the responding cells were from the same young donor.

**Figure 4.**
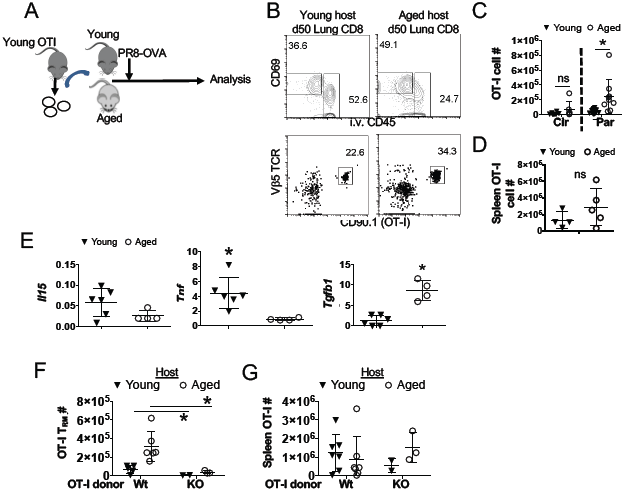
Excessive accumulation of T_RM_ cells in an aged environment is dependent on TGF-βR signaling. **A.** OT-I T cells (CD90.1^+^) from young donors were adoptively transferred into young or aged C57BL/6 mice 1 day prior to infection with PR8-OVA virus. **B.** Representative flow cytometry plots in lungs at 50 d.p.i., showing the gating strategy for resident OT-I T cells (CD8^+^CD69^+^ i.v.CD45^-^ Vβ5^+^CD90.1^+^) **C, D.** Donor OT-I cells were enumerated in young and aged hosts in the lung vasculature (Cir) or parenchyma (Res) (C), or in spleens (D) at 50 d.p.i. **E.** Relative transcripts (RT) of *Il-15, Tnf*, and *Tgfb1* (left to right) from total lung samples obtained from aged and young animals at 60 d.p.i. was evaluated by qPCR. **F, G.** Wild type (WT, CD45.1^+^) or *TGFbR2*^*fl/fl*^ *dLck-Cre* OT-I cells (KO, CD45.1^+^) were adoptively transferred from young donors into young or aged hosts (CD45.2^+^) one day prior to PR8-OVA infection. **F.** Resident OT-I cells were enumerated in the lung of young or aged mice at 50 d.p.i. **G.** OT-I cells in the spleen of young or aged mice at 50 d.p.i. **C** is pooled data of 3 independent experiments, **E-G.** are representatives of 2 repeats. * p<0.05 Student’s two-tailed t-test with unequal variance.

To explore the potential mechanisms regulating CD8^+^ T_RM_ cell accumulation during aging, we performed qRT-PCR analysis on infected lungs to examine the expression of *Il15, Tnf* and *Tgfb1*, which are important in T_RM_ cell development and/or maintenance [36-40]. We found that *Tgfb1* transcript levels were elevated in infected aged lungs compared to those of infected young lungs (Figure 4E). Previously, TGF-βR signaling was shown to be critical for the development of both CD69^+^ CD103^+^ and CD69^+^ CD103^-^ lung T_RM_ cells [41]. Therefore, we examined whether the excessive accumulation of CD8^+^ T_RM_ cells in aged lungs is dependent on TGF-βR signaling. To this end, we adoptively transferred young WT or TGFβRII-deficient (TGFβRII KO, dlLck-Cre Tgfbr2^fl/fl^). OT-I cells into young or aged hosts, and then infected the mice with PR8-OVA. TGFβRII deficiency not only diminished OT-I T_RM_ development in young mice, but also abolished the excessive accumulation of OT-I T_RM_ in aged mice (Figure 4F). However, TGFβRII deficiency did not significantly affect splenic memory T cell responses in young or aged mice (Figure 4G). Thus, these data suggest that excessive accumulation of CD8^+^ T_RM_ cells in aged lungs is dependent on d TGF-βR signals in aged tissue environment.

### Age-associated dysfunction of T_RM_ cells against a major protective epitope

T_RM_ cells are vital for heterologous influenza virus reinfection [43]. Since aged lungs accumulated significant more CD8^+^ T_RM_ cells, it is plausible that aged mice may exhibit enhanced protective heterologous immunity against viral reinfection. To this end, we employed a heterologous infection and challenge model in which we infected young or aged mice with PR8 virus and then re-challenge the mice with a lethal dose of X31 virus in the presence of FTY720, which blocks the contribution of protective circulating memory T cells [12]. PR8 and X31 viruses differ in viral surface proteins, but share internal viral proteins such as NP, which are cross-recognized by memory CD8^+^ T cells [44, 45]. We confirmed that FTY720 treatment abolished T cell migration (Figure S4A). We then followed host morbidity and mortality daily following X31 challenge. All naïve young or aged mice succumbed to lethal X31 infection (Figure S4B), while PR8-infected young mice were fully protected from lethal X31 infection in the presence of FTY720 (Figure 5A). Strikingly, ∼75% of PR8-infected aged animals succumbed to secondary X31 re-challenge, which was almost comparable to mortality of primary X31 infection in those mice (Figure 5A and S4B). Thus, the enhanced presence of CD8^+^ T_RM_ cells in aged lungs failed to provide heterologous protection against influenza reinfection.

**Figure 5.**
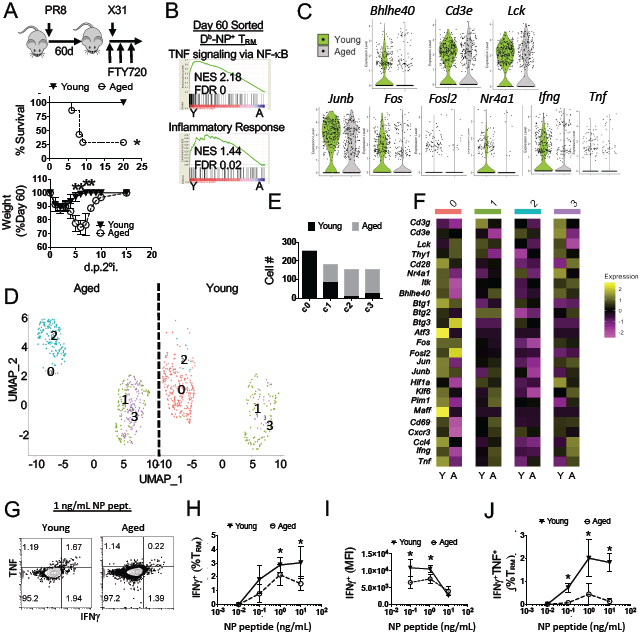
D^b^-NP T_RM_ cells from aged lungs are dysfunctional in protective immunity. **A.** PR8 virus infected young (filled triangles, Y) or aged (open circles, A) mice were treated with FTY720 daily starting at 60 d.p.i. per methods and materials. Then mice were infected with a lethal dose of X31 at 61 d.p.i. Survival (middle panel) and weight loss (bottom) as percent of pre-secondary infection weight were determined daily post secondary infection (d.p.2°i.). **B.** D^b^-NP-specific T_RM_ cells (CD8^+^CD44^Hi^CD69^+^i.v.CD90^-^D^b^-NP tetramer^+^) from PR8 infected young (n=18) or aged (n=12) mice were pooled and sorted for single-cell RNAseq analysis. GSEA of NF-кB signaling and inflammatory response. NES: normalized enrichment score, FDR: False discovery rate (middle and bottom panel). **C.** Violin plots of *Bhlhe40, Cd3e, Lck, Junb, Fos, Fosl2, Nr4a1, Ifng* and *Tnf* expression in young or aged D^b^-NP T_RM_ cells. **D.** Seurat UMAP plots with clusters 0-3 of aged (left panel) and young (right panel) samples from scRNAseq data. **E.** Composition of aged or young T_RM_ cells per cluster. **F.** Cluster-differentiated heat map (purple = lower relative expression, yellow = higher relative expression) of indicated genes. **G.** Representative flow cytometry plots of the production of IFN-γ and TNF in lung resident CD8^+^ T cells following 1 ng/ml NP 366-374 peptide stimulation. **H.** Percent of IFN-γ^+^ resident CD8^+^ T cells following stimulation with increasing amount of NP 366-374 peptide (0.01 – 10 ng/mL). **I.** Geometric Mean Fluorescence Intensity (gMFI) IFN-γ levels following of stimulation with increasing amount of NP 366-374 peptide. **J.** Percent of IFN-γ^+^ TNF^+^ resident CD8^+^ T cells following stimulation with increasing amount of NP 366-374 peptide (0.01 – 10 ng/mL). **A** are pooled data from 2 experiments. **G-J** are representatives of 3 experiments. * p<0.05 Student’s two-tailed t-test with unequal variance.

Among influenza-specific CD8^+^ T cells, D^b^-NP-specific T cells appear to be particularly critical for the protection against secondary heterologous viruses [46]. To examine the mechanisms underlying the impaired protective function of T_RM_ cells in aged lungs, we sorted polyclonal D^b^-NP-specific T_RM_ from infected young or aged lungs, and performed scRNAseq at 60 d.p.i. (Figure S4C). Compared to D^b^-NP T_RM_ cells from aged mice, D^b^-NP T_RM_ from the young mice were enriched in genesets associated with NF-кB signaling and inflammatory immune responses (Figure 5B). Furthermore, D^b^-NP T_RM_ from aged mice had decreased expression of effector molecules such as *Ifng* and *Tnf*, and molecules involved with TCR signaling such as *Lck* and *Junb* (Figure 5C). Additionally, *Bhlhe40*, a key transcription factor necessary for the expression of effector molecules in T_RM_ cells [13], was diminished in T_RM_ from aged lungs (Figure 5C). We performed Uniform Manifold Approximation and Projection (UMAP) and found that D^b^-NP T_RM_ from young and aged mice can be divided into 4 clusters (Figure 5D and S4D). Cluster 0 cells were mainly comprised of T_RM_ cells from young mice, and clusters 2 and 3 cells consisted mainly of T_RM_ cells from aged mice, while cluster 1 cells were equally divided by T_RM_ cells from young and aged mice (Figure 5D & E). Compared to the rest of cluster cells, cells in cluster 0 express high levels of genes associated with classical T_RM_ signature (such as *Bhleh40, Cd69* and *Jun*), molecules associated with better TCR signaling (such as *Lck, Cd3e* and *Cd28*) and genes ^+^ T cell effector function (such as *Ccl4* and *Tnf*) (Figure 5F). Thus, associated with CD8 age-associated lung T_RM_ cells lose a functional subpopulation that is characterized with robust expression of CD8^+^ T cell effector and functional molecules, which likely explain the lack of protection against secondary viral re-challenge.

To further test the capacity of T_RM_ to elicit a protective response, we stimulated lung cells from young or aged mice with NP 366-374 peptide and then measured IFN-γ and TNF production by CD8^+^ T_RM_ cells through intracellular staining. Consistent with the scRNAseq data, D^b^-NP-specific T_RM_ cells from aged mice exhibited diminished production of IFN-γ compared to young mice, particularly on the IFN-γ levels were evaluated on a per cell basis (Figure 5 G-I). Moreover, CD8^+^ T_RM_ cells from aged lungs were defective in simultaneously producing both IFN-γ and TNF following peptide re-stimulation (Figure 5J). Of note, splenic CD8^+^ memory T cells were also impaired in producing both IFN-γ and TNF following peptide re-stimulation during aging (Figure S5A), suggesting that the impaired production of functional effector molecules following peptide re-stimulation was not restricted to T_RM_ cells. In contrast to their diminished cytokine production following peptide stimulation (TCR signaling), total CD8^+^ or D^b^-NP-specific T_RM_ cells were capable of producing IFN-γ and TNF following PMA/Ionomycin restimulation (Figure S5B, C). These data suggest that D^b^-NP T_RM_ cells exhibit impaired production of effector molecules specifically following TCR stimulation. Interestingly, following ex vivo stimulation of D^b^-PA specific T_RM_ cells by PA 224-233 peptide, we found that D^b^-PA T_RM_ cells from aged and young hosts had similar ability to produce IFN-γ and TNF (Figure S5D). These data suggest that the impaired production of effector molecules by D^b^-NP-specific T_RM_ cells is likely not due to the aged environment, but their developmental defects as demonstrated before [28]. Consistent with this idea, young OT-I cells transferred into the aged hosts did not exhibit functional defects in cytokine production compared to OT-I cells transferred into young mice (Figure S5 E & F).

### Resident CD8^+^ T cells in aged hosts drive persistent pulmonary inflammation and lung fibrosis

Excessive accumulation of T_RM_ cells following PD-L1 blockade in influenza-infected mice leads to persistent inflammatory and fibrotic responses in the lungs at the memory stage [41]. Since aged lungs harbored enhanced T_RM_ cells and persistent lung pathology, we hypothesized that excessive accumulation of CD8^+^ T_RM_ in aged lungs may cause persistent inflammation and fibrotic sequela. To test this hypothesis, we infected aged mice with PR8 virus and then injected the mice with high or low doses of CD8 depleting antibody (α-CD8) at 21 d.p.i. to deplete CD8^+^ T cells systemically in lymphoid and peripheral tissues (high dose) or only in lymphoid organs and the circulation (low dose) [40, 47, 48] (Figure 6A). Both low dose and high dose CD8 Ab treatment largely ablated splenic CD8^+^ T cells, however, only high dose CD8 Ab injection caused significant ablation of CD8^+^ T cells in the lung parenchyma (Figure 6B and S6A). Neither high nor low dose of CD8 Ab treatment caused significant reduction of CD4^+^ T or B cell compartments in the lungs (Figure S6 B). Lung histopathological analysis revealed marked reduction of inflammatory lesions following high dose, but not low dose of CD8 Ab treatment particularly in aged mice following infection (Figure 6 C-E), indicating that lung resident CD8^+^ T cells drive chronic lung pathology at the memory phase. Consistent with the histology data, high dose, but not low dose, of CD8 Ab treatment caused significant reduction of Ly6C^+^ monocyte and neutrophil infiltration in aged, but not young lung tissues (Figure 6F). Thus, resident CD8^+^ T cells are responsible for the constant recruitment of monocytes and neutrophils to the tissue at the memory phase. To further explore the roles of resident CD8 T cells in promoting tissue inflammation at the memory stage, we measured immune-associated genes in the lungs by Nanostring. There was a marked decrease in the expression of multiple cytokines and particular chemokines in the lungs of mice that received high CD8 Ab treatment, compared to those of lungs of mice received control Ab or low dose of CD8 Ab at 60 d.p.i. (Figure 6G & 6SC & D), likely explaining the diminished monocyte and neutrophil infiltration in the tissue following resident CD8^+^ T cell depletion.

**Figure 6.**
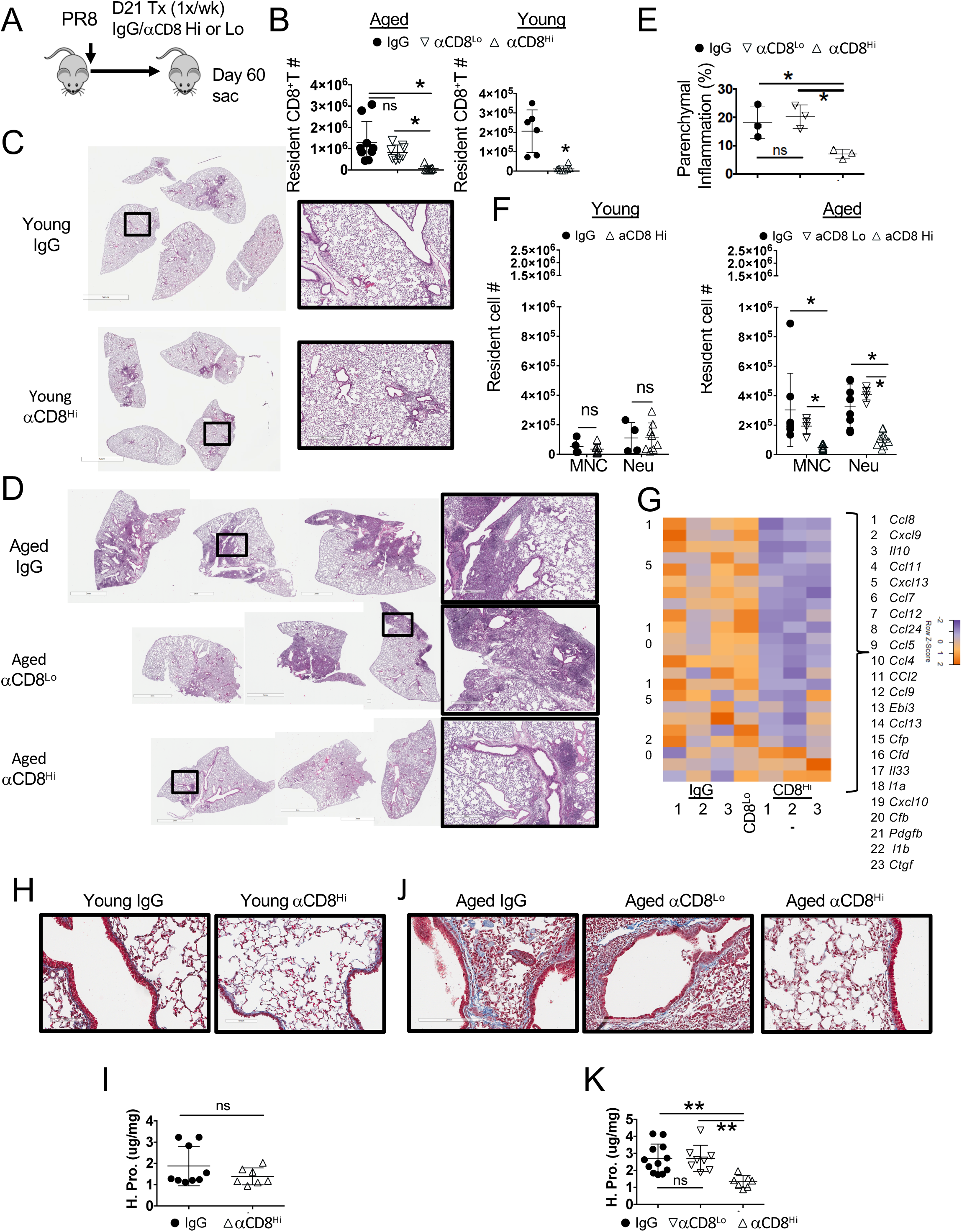
Resident CD8^+^ T cells support chronic parenchymal inflammation and fibrosis in aged hosts. Aged or young WT mice were infected with PR8 strain. **A.** Experimental procedure of high or low dose of CD8 antibody (αCD8) treatment starting at day 21 post-infection. **B.** Lung-resident CD8^+^ T cell numbers in aged (left panel) or young (right panel) mice following CD8 Ab treatment. **C, D.** H&E staining of whole left-lung lobes from young **(C)** or aged mice (**D**) treated with indicated Abs. **E.** Percent of left lung lobes infiltrated by white blood cells via Image J analysis of D. **F.** Quantitation of resident monocytes (Ly6C Hi, ivCD45^-^CD64^-^CD11c^Lo^Siglec-F^-^ CD11b^Hi^Ly6C^Hi or Lo^) and neutrophils (Neu, ivCD45^-^Ly6G^Hi^CD11b^Hi^) in IgG or high dose CD8 depleted young (left) or aged (right) mice treated with indicated Ab. **G.** Heat map for select DEGs of the lungs from aged mice treated with IgG (in triplicate), high dose (αCD8^Hi^, in triplicate) or low dose (αCD8^Lo^, pooled n=3) of CD8 Ab as analyzed by Nanostring. Orange is relatively high expression and purple is relatively low expression. **H.** Representative 200x micrographs of Mason’s trichromatic stain on lung sections of young mice with or without high dose CD8 Ab treatment. **I.** Hydroxyproline levels reflective collagen content of the lungs from young mice with or without high dose CD8 Ab treatment. **J.** Representative 200x micrographs of Mason’s trichromatic stain on lung sections of aged mice received control Ab, low dose of CD8 Ab (αCD8^Lo^) or high dose CD8 Ab (αCD8^Hi^) treatment. **K.** Hydroxyproline levels reflective collagen content of the lungs from aged mice received control Ab, low dose of CD8 Ab (αCD8^Lo^) or high dose CD8 Ab (αCD8^Hi^) treatment. **B, F, H-K** are 2 pooled experiments, **E & G** are single replicates. * p<0.05 Student’s two-tailed t-test with unequal variance **(I)** or for ANOVA with correction for multiple tests **(B, E, F, J, & K)**.

Acute influenza virus infection can cause persistent lung remodeling and collagen deposition [24]. We therefore examined whether CD8^+^ T cell depletion at 3 weeks post infection could affect lung fibrosis at the memory stage. CD8^+^ T cell depletion did not affect lung collagen content in infected young mice revealed by both Mason’s trichrome staining and hydroxyproline assay (Figure 6H and I), which measures total collagen protein [49]. These data suggest that normal CD8^+^ T_RM_ cell responses in young mice contribute minimally to lung fibrotic responses following acute influenza virus infection, which is in contrast to the excessive CD8^+^ T_RM_ cell-induced fibrotic sequelae following PD-L1 blockade [41]. In contrast to young mice, aged mice exhibited increased lung collagen deposition, particularly in the lung parenchyma (Figure 6H & J). Strikingly, mice received high CD8 Ab treatment had markedly reduced lung fibrosis as revealed by Trichome staining and hydroxyproline assay, compared to lungs of mice that received control Ab or low dose of CD8 Ab (Figure 6J & K). These data suggest that resident but not circulating CD8^+^ T cells are important in promoting chronic lung collagen deposition following the resolution of primary influenza infection. In summary, our data indicate that excessive accumulation of lung resident CD8^+^ T cells does not lead to enhanced protective immunity in aged mice, but rather drive age-associated persistent lung inflammation and chronic fibrosis following viral pneumonia.

## Discussion

In this report, we demonstrate that acute influenza infection leads to persistent chronic pulmonary inflammation and fibrosis, which are largely propagated by an exaggerated CD8^+^ T_RM_ response in aged hosts. Somewhat counterintuitively, elevated CD8^+^ T_RM_ cells in the aged lung, fail to provide the protective heterologous immunity to that observed in young mice, likely due to a selective dysfunction in a protective CD8^+^ T_RM_ cell population (i.e. D^b^-NP CD8 T cells) [50, 51].

Previous reports suggest that both T cell-intrinsic and extrinsic environmental factors could contribute to the suboptimal effector and memory CD8^+^ T cell responses during aging [8, 52, 53]. To our surprise, we observed enhanced T_RM_ cell accumulation, despite diminished lung effector T cell responses and lower levels of splenic memory T cells observed in aged hosts. Our data further indicate that aged environment, likely through elevated tissue TGF-β, promotes T_RM_ cell generation and/or maintenance in a T cell-extrinsic fashion during aging. Of note, aging has been linked to altered naïve T cell repetoires and enhanced representation of senescent virtual memory T cells [54]. Whether these T cell intrinsic changes in aged hosts, in addition to environmental factors, contribute to an enhanced T_RM_ generation, requires further investigation. Notably, despite a numeric increase in T_RM_ cells, protective heterologous immunity is impaired in aged hosts due to a qualitative defect in T_RM_ cells not present in young mice. The protective function of T_RM_ cells is usually associated with their effector activities at the site of pathogen re-entry [56]. For instance, it has been shown that IFN-γ is important for T_RM_ cell protective immunity against secondary heterologous influenza virus infection [43]. Consistent with the notion, scRNAseq analysis found that T_RM_ cells against a major protective CD8^+^ T cell epitope (D^b^-NP) from aged mice lose a functional T_RM_ subset observed in young hosts; this subset was characterized with robust expression of molecules associated with TCR signaling and effector molecules. As a result, D^b^-NP specific T_RM_ cells are less sensitive to TCR stimulation for the simultaneous production of IFN-γ and TNF. Thus, defective TCR-mediated effector cytokine production could underlie the impaired protection against secondary influenza virus infection. Notably, T_RM_ cells can undergo *in situ* proliferation and generate secondary effector T cells [41, 57, 58]. Previously, it has been shown that memory T cells in aged mice are senescent in proliferation during secondary expansion [8]. Therefore, it is also possible that impaired T_RM_ proliferation and secondary effector T cell expansion could together contribute to the impaired T_RM_ protective function.

Interestingly, impaired effector cytokine production by D^b^-NP T_RM_ cells was not observed in D^b^-PA or transferred young OT-I T_RM_ cells, suggesting that this functional defect in D^b^-NP T cells is likely not caused by the aged environment. It has been shown that D^b^-NP, but not D^b^-PA, CD8^+^ T cells have preferential age-associated loss in responsiveness, which is mainly due to the narrowed TCR-β repertoire in D^b^-NP CD8^+^ T cells or their recruitment into a response [6, 28, 32]. Thus, intrinsic developmental defects are likely the cause of the functional decline in D^b^-NP T_RM_ cells [28]. Notably, our data highlight the importance of D^b^-NP T_RM_ cells in protective immunity as functional impairment specifically in D^b^-NP cells could translate into differences in the secondary protective immunity. This notion is consistent with previous reports, which have pinpointed robust expansion of D^b^-NP memory T cells over D^b^-PA memory T cells in the secondary expansion to provide enhanced protective immunity following heterologous challenge [46, 59].

CD8^+^ T_RM_ cells exhibit higher levels of expression of multiple effector molecules compared to central and effector memory T cells [13, 60]. The heightened expression of these molecules confers CD8^+^ T_RM_ cells with superb antiviral function, but may potentially cause bystander tissue inflammation and injury [41]. In experimental models, both CD4^+^ and CD8^+^ T_RM_ cells have been implicated in causing tissue immunopathology ranging from persistent allergic inflammation to inflammatory bowel disease [61, 62]. Previous reports have demonstrated that influenza infection in young mice leads to chronic inflammation and collagen deposition [24]. Notably, resident CD8^+^ T cell depletion only moderately affected pulmonary inflammation and lung collagen deposition in young mice, even though exaggerated CD8^+^ T_RM_ responses following PD-1 blockade can lead to inflammatory and fibrotic sequelae [41]. These data suggest that influenza-infected young lungs are able to tolerate “normal levels” of T_RM_ cells, likely due to the expression of various inhibitory molecules expressed by T_RM_ cells [41]. During aging, however, influenza virus infection leads to excessive accumulation of T_RM_ cells, which exceeds the ability of the aged tissue to “counterbalance” the pathogenic activities of T_RM_ cells, driving enhanced pulmonary inflammation and parenchymal fibrosis. The cellular and molecular mechanisms underlying T_RM_-mediated lung pathology and fibrosis during aging are unclear currently. However, aging is associated with the impaired capability of tissue repair [27]. It is possible that epithelial injury caused by persistent low level release of cytopathic molecules by T_RM_ cells surpasses the limited recovery capability of the aged tissue. To this end, even though D^b^-NP T_RM_ cells have diminished TCR signaling for IFN-γ and TNF production, their responses to PMA and Ionomycin are intact. Therefore, they may remain capable of producing low levels of effector molecules in response to environmental inflammatory factors that are constitutively observed in aged tissues [22, 27]. Alternatively, it is possible that enhanced accumulation of those “functionally-intact” T_RM_ cells such as D^b^-PA T_RM_ cells may be responsible for the persistent tissue inflammation and fibrosis in aged hosts. Additionally, T_RM_ cells in aged mice may have different traits than T_RM_ cells in young mice [63, 64], thereby causing increased lung inflammation and tissue fibrosis.

Aging is associated with enhanced development of a number of chronic lung diseases including chronic obstructive pulmonary diseases (COPD) and pulmonary fibrosis [65]. Interestingly, aging is also associated with increased susceptibility of acute lung injury and severe disease development following respiratory viral infections such as influenza virus and SARS-CoV-2 [17, 66, 67]. Increasing evidence has suggested that acute viral infections may cause persistent lung function impairments due to pulmonary fibrosis [20]. Indeed, lung fibrosis is well-documented in a substantial number of patients who have recovered from the infection of SARS-CoV or MERS-CoV [68, 69] and a group of influenza patients with severe acute diseases [20]. Emerging evidence has also suggested that COVID-19 survivors will likely exhibit persistent impairment of lung function and the development of pulmonary fibrosis [21, 70-74]. Data presented here suggests for the first time that excessive T_RM_ cells are likely the cause of chronic lung diseases and pulmonary fibrosis following viral pneumonia in aged individuals. Of note, recent data on scRNAseq analyses have suggested that pulmonary T cells from COVID-19 patients are enriched with T_RM_ signature and CD8^+^ T cell clonal expansion in the airways (doi: https://doi.org/10.1101/2020.02.23.20026690). Thus, strategies aimed at decreasing exuberant T_RM_ responses and/or inhibiting the pathogenic potential of T_RM_ cells may be promising in treating chronic lung diseases following viral pneumonia in the elderly, including COVID-19 survivors.

In summary, we have unveiled an age-associated paradox in CD8^+^ T_RM_ cell responses. The excessive accumulation of T_RM_ cells is not protective but rather drives inflammatory and fibrotic sequalae following primary respiratory viral infection. Moving forward, it is crucial to better understand the mechanisms of age-associated T_RM_ malfunction so that we can selectively restore T_RM_ protective function while minimizing their pathogenic potential in the elderly.

## Materials & Methods

### Mice, infections, and adoptive transfers

C57BL/6 were originally purchased from Jackson (Harbor, ME) Laboratory and mostly bred in house. Aged mice were received at 20-21 months of age from the National Institutes of Aging and maintained in same specific-pathogen-free conditions for at least one month prior to infection. In most of cases, aged and young subject bedding was cross-contaminated weekly to control for different microbial environments. Young OT-I and dLck-Cre Tgfbr2^fl/fl^ OT-I lymphoid tissues (KO, CD45.1^+^) were shipped from Nu Zhang (University of TX Health Science Center at San Antonio). All mice were housed in a specific pathogen-free environment and used under conditions fully reviewed and approved by the internal animal care and use committee (IACUC) guidelines at the Mayo Clinic (Rochester, MN). For primary influenza virus infection, influenza A/PR8/34 strain (∼100 pfu/mouse) was diluted in FBS-free DMEM media (Corning) on ice and inoculated in anesthetized mice through intranasal route as described before [76]. X-31 strain was prepared identically and 2.8×10^5^ pfu was administered per mouse for secondary challenge. During the secondary challenge phase, experimental mice were treated with 1 μg/g FTY720 daily starting one day prior to re-challenge.. For adoptive transfers, 1×10^4^ OT-I from lymph nodes of 8-10 weeks old females were transferred intravenously and mice were infected as indicated 1 day later.

### Cell depletions

For depletion of CD8 T cells, starting at day 21 post-infection, mice were given 20 μg (Low dose) or 500 μg of anti-CD8 Ab (53-6.7) or control rat IgG (500 μg) in 200 μL PBS through once weekly intraperitoneal injections. Mice were sacrificed 3 days after the last treatment.

### Parabiosis surgery

To examine tissue residency of lung CD69^+^ parenchymal CD8^+^ T cells, parabiotic surgery was performed. CD45.1^+^ young naive mice were paired with infected aged CD45.2^+^ congenic mice at 35 d.p.i. Briefly, mice were anesthetized with ketamine and xylazine. After disinfection, incisions were made in the shaved skin area then the olecranon of the knee joints of each mouse was joined. Opposing incisions were closed with a continuous suture on the dorsal and ventral sides. Parabionts were then allowed to rest for 4 weeks before sacrifice and examining the lung vasculature, parenchyma, and spleen immuno-profiles. Equilibration of white blood cells between parabionts was confirmed in the peripheral blood before tissue analysis by flow cytometry.

### Tissue Processing, cellular isolation, stimulations, and data analysis

Animals were injected intravenously with 4 μg of CD45 or 2 μg CD90.2 antibody labeled with various fluorochromes. 2 minutes post-injection, animals were euthanized with an overdose of ketamine/xylazine and processed 3 minutes later. Following euthanasia, spleens were removed and the right ventrical of the heart was gently perfused with PBS (10 mL). Lungs were instilled with either 1 mL of 10% formalin for histology studies or 1 mL of digestion buffer (90%DMEM 10% PBS+Calcium with 180 U/mL Type 2 Collagenase (Worthington) and 15 μg/mL DNase (Sigma) additives). Tissue was processed on a gentleMACS tissue disrupter (Miltenyi) for 40 minutes at 37°C followed by hypotonic lysis of red blood cells in ammonium-chloride-potassium buffer and filtering through 70 μm mesh. FC-gamma receptors were blocked with anti CD16/32 (2.4G2). Cells were washed twice in PBS and stained with near infrared Zombie viability dye per manufacturers protocol (Biolegend). Cell surfaces were immuno-stained with the following cocktails of fluorochrome-conjugated Abs (Biolegend). Immuno-staining was performed at 4 °C for 30 min. Cells were washed twice with FACS buffer (PBS, 2 mM EDTA, 2% FBS, 0.09% Sodium Azide), prior to fixation and ran on an Attune NxT autosampler (Life Technologies). FCS files for myeloid stains were analyzed with FlowJo 10.2 (Tree Star). Antibody clones are as following: Siglec-F (E50-2440), CD11c (N418), CD11b (M1/70), CD64 (X54-4/7.1), Ly6G (1A8), I-A/I-E (M5/114.15.2), Ly6C (HK1.4), CD8α (53-6.7), CD69 (H1.2F3), CD44 (IM7), CD103 (2E7), PD-1 (29F.1A12). For functional assays, following digestion protocols above, cells were resuspended at 3×10^6^/mL in RPMI with 10% FCS and stimulated with indicated amount of peptide or PMA and Ionomycin for 3 hours before adding monensin for the last 2 hours of culture.

### Hydroxyproline assay

50 mg lung tissue was hydrolyzed in 1 ml of 6 M HCl at 95°C overnight. Hydroxyproline standard solution was purchased from Sigma-Aldrich. Hydroxyproline concentration is determined by the reaction of oxidized hydroxyproline with 4-(Dimethylamino)benzaldehyde (DMAB). The product was read at 560-nm wavelength in a Thermax plate reader as described [41, 77].

### Quantification of Histopathology

Hematoxylin and eosin stained formalin-fixed lung sections were scanned using an Aperio Image Scanner (leica). Tiff files were converted to 16-bit black and white and two threshold measurements were taken using Image J software (NIH). One measured the total parenchymal area and the other the inflamed area of the tissue. The caveat of the method is that bronchiole epithelium is thresholded in both measurements. To account for this, values from a group of age-matched naive mice were subtracted from the above ratios to yield % of inflamed/infiltrated parenchyma.

### Quantitative RT-PCR

Total RNA was extracted from lung tissue from Trizol preparations. Random primers (Invitrogen) and MMLV reverse transcriptase (Invitrogen) were used to synthesize first-strand cDNAs from equivalent amounts of RNA from each sample. These cDNA were subjected to realtime-PCR with Fast SYBR Green PCR Master Mix (Applied Biosystems). Realtime-PCR was conducted on QuantStudio3 (AppliedBioscience). Data were generated with the comparative threshold cycle (Delta CT) method by normalizing to hypoxanthine phosphoribosyltransferase (HPRT) transcripts in each sample as reported previously [78].

### Nanostring analysis

Total RNA was extracted from bulk lung samples in Trizol 0 or 60 d.p.i. 100 ng of RNA was either pooled between group or used from individual samples as indicated. Library hybridization was established by following the instructions of the manufacturer. Aliquots of Reporter CodeSet and Capture ProbeSet were thawed at RT. Then, a master mix was created by adding 70 µl of hybridization buffer to the tube containing the reporter codeset. Eight microliters of this master mix was added to each of the tubes for different samples; 5 µl of the total RNA sample was added into each tube. Then, 2 µl of the well-mixed Capture probeset was added to each tube and placed in the preheated 65°C thermal cycler. All the sample mixes were incubated for 18 hours at 65°C for completion of hybridization. The samples were then loaded into the sample hole in the cartridge and loaded into the NanoString nCounter SPRINT Profiler machine (NanoString). When the corresponding Reporter Library File (RLF) running was finished, the raw data were downloaded and analyzed with NanoString Software nSolver 3.0 (NanoString). mRNA counts were processed to account for hybridization efficiency, background noise, and sample content, and were normalized using the geometric mean of housekeeping genes. All data were normalized by housekeeping genes. Heat maps were generated by an assortment of R language packages.

### Single-cell RNA sequencing

Sorted (CD8^+^CD44^Hi^CD69^+^i.v.CD90^-^D^b^-NP tetramer PE/APC-double fluorochrome positive) T cells from pooled lung cells of mice (10-18 mice) infected with influenza virus (60 d.p.i.) were loaded on the Chromium Controller (10x Genomics). Single-cell libraries were prepared using the Chromium Single Cell 3’ Reagent kit (10x Genomics) following manufacturer’s instruction. Paired-end sequencing was performed using an Illumina HiSeq 2500 in rapid-run mode. CellRanger software package (10x Genomics) were used to align and quantify sequencing data from 10x Genomics. All scRNA-seq analyses were performed in R using the Seurat package (version 3.0).

### Total RNA-sequencing

RNA was extracted from bulk lung samples with trizol. After quality control, high quality (Agilent Bioanalyzer RIN >7.0) total RNA was used to generate the RNA sequencing library. cDNA synthesis, end-repair, A-base addition, and ligation of the Illumina indexed adapters were performed according to the TruSeq RNA Sample Prep Kit v2 (Illumina, San Diego, CA). The concentration and size distribution of the completed libraries was determined using an Agilent Bioanalyzer DNA 1000 chip (Santa Clara, CA) and Qubit fluorometry (Invitrogen, Carlsbad, CA). Paired-end libraries were sequenced on an Illumina HiSeq 4000 following Illumina’s standard protocol using the Illumina cBot and HiSeq 3000/4000 PE Cluster Kit. Base-calling was performed using Illumina’s RTA software (version 2.5.2). Paired-end RNA-seq reads were aligned to the mouse reference genome (GRCm38/mm10) using RNA-seq spliced read mapper Tophat2 (v2.1.1). Pre- and post-alignment quality controls, gene level raw read count and normalized read count (i.e. FPKM) were performed using RSeQC package (v2.3.6) with NCBI mouse RefSeq gene model. For functional analysis, GSEA was used to identify enriched gene sets, from the hallmark and C5 databases of MSigDB, having up-regulated and down-regulated genes, using a weighted enrichment statistic and a log2 ratio metric for ranking genes.

### Statistical analysis

Quantitative data are presented as mean ± Standard of Deviation. Unpaired two-tailed Student’s t-test (two-tailed, unequal variance) was used to determine statistical significance with Prism software (Graphpad). Where appropriate, ANOVA corrected for multiple comparisons was used (Graphpad). We considered *p* < 0.05 as significant in all statistical tests and denoted within figures as a * for each order of magnitude *p* value.

## Acknowledgements

We thank NIH tetramer core for tetramers and Mayo flow cytometry core and genomic core for technical assistance. This study was funded by the US National Institutes of Health grants AI112844, AI147394 and AG047156 to J.S., NS103212 to A.J., AI125701 and AI139721 to N.Z. and HL52160 to A.L. Mayo Clinic CBD Research Fund to J.S., E.C. and R.V. Cancer Research Institute CLIP program and American Cancer Society grant RSG-18-222-01-LIB to N.Z. Three Lakes Partners Foundation to A.L.

---

**Figure S1.**
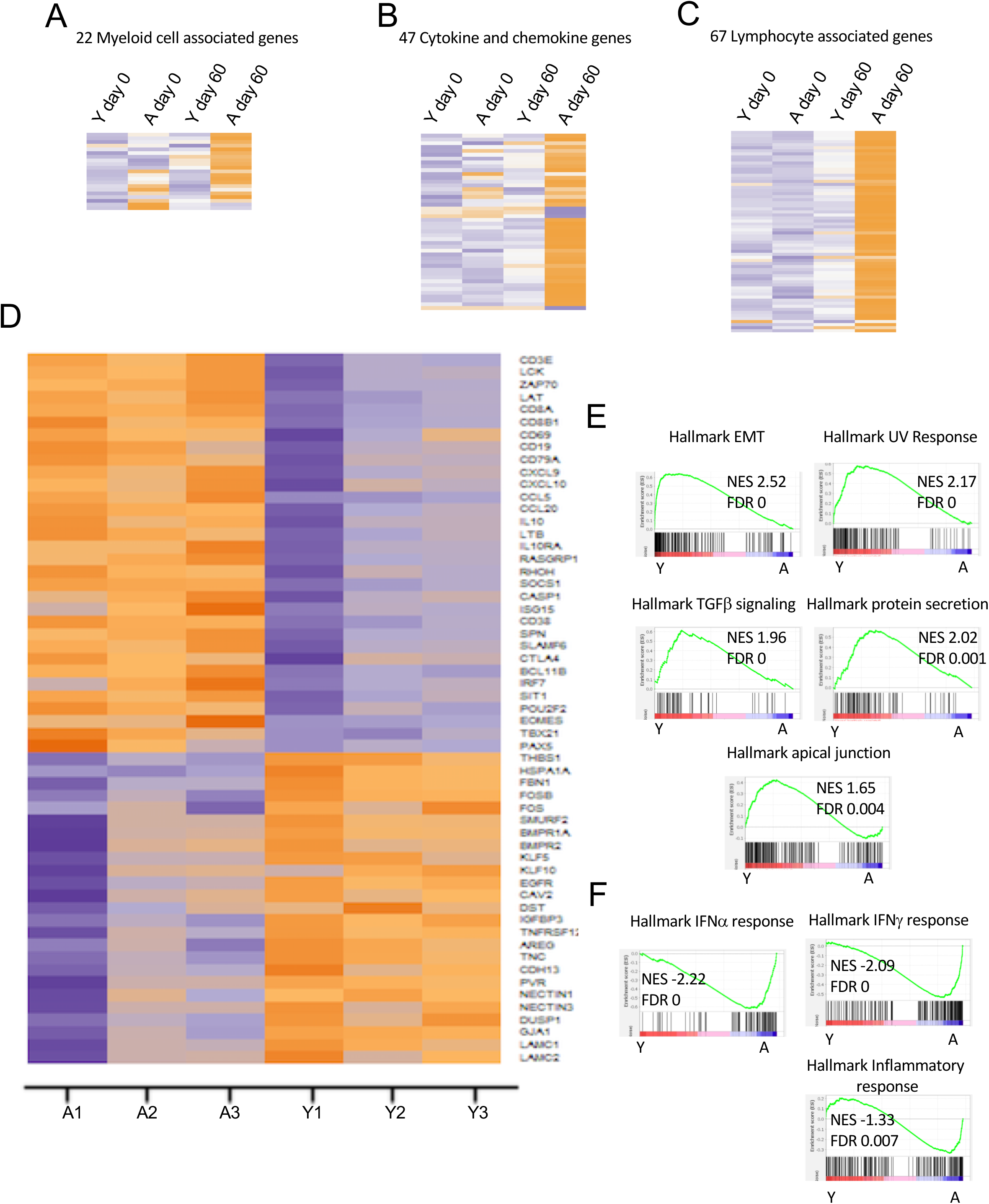
Aged lungs exhibit persistent inflammatory responses following influenza infection. **A-C.** Young (Y) or aged (A) C57BL/6 mice were infected (day 60) or not (day 0) with PR8. 560 immune-associated genes in the lungs were analyzed by Nanostring (at least 3 pooled samples/group). Myeloid **(A)**, cytokine and chemokine (**B)**, or lymphocyte (**C)** associated DEGs that were at least 1.5 fold expression level changes from Aged to young infected samples. **(D)** RNAseq heatmap showing DEGs in young or aged infected lungs 60 d.p.i. **E, F.** GSEA plots of RNAseq data showing genes enriched in young infected lungs **(D)** or aged infected lungs (**F)**. NES: normalized enrichment scores and FDR: false-discovery rates. Nanostring data is representative of 2 experiments with pooled samples. RNAseq data is from a single replicate with samples in triplicate.

**Figure S2.**
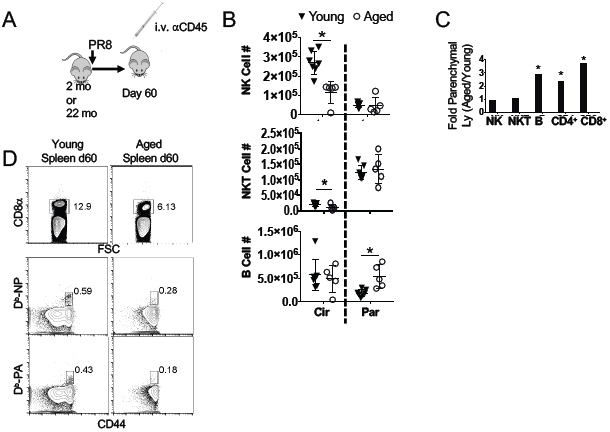
Enhanced presence of adaptive immune cells in aged lung parenchyma. Young (filled triangles) or aged (open circles) C57BL/6 mice were infected with PR8 and injected intravenously (i.v.) with anti-CD45 to label circulating white blood cells prior to sacrifice. **A.** Schematics of experimental procedure. **B.** NK, NKT, and B cells were enumerated in the lung vasculature (Cir; left panel) and parenchyma (Par; right panel) at 60 d.p.i. **C.** Fold change of aged/young cells in parenchyma with indicated lymphocyte population at 60 d.p.i. **E.** Representative flow cytometry gating of CD8^+^ CD44^Hi^ D^b^-tetramer^+^ NP and PA specific memory T cells in the spleen at 60 d.p.i. B-D are representatives o**f** 3 experiments. * p<0.05 Student’s two-tailed t-test with unequal variance.

**Figure S3.**
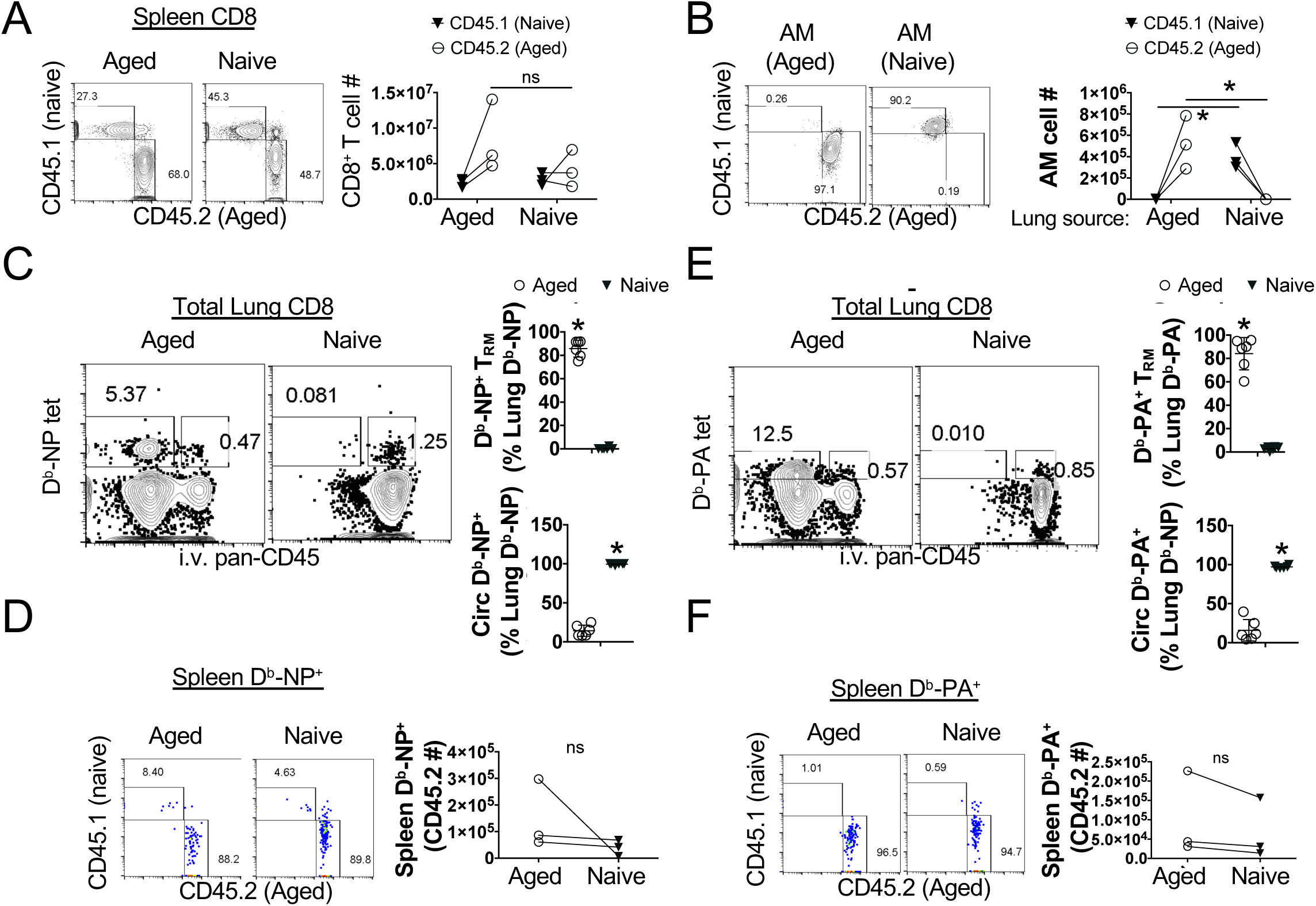
Aged parenchymal CD8^+^ CD69^+^ memory T cells are tissue resident. Aged mice (CD45.2^+^) were infected with PR8 and parabiosed with young naive (CD45.1^+^) mice at 5 weeks after infection. **A.** Representative flow cytometry plots (left panel) or cell numbers (right panel) of splenic CD8^+^ T cells, showing the host or donor derived cells in infected aged or naive hosts. **B.** Representative flow cytometry plots (left panel) or cell numbers (right panel) of alveolar macrophages, showing the host or donor derived cells in infected aged or naive hosts. **C.** Representative flow plot of total lung CD8 T cells showing D^b^-NP^+^ tetramer staining in the parenchyma (CD45 i.v.^-^) or circulation (CD45 i.v.^+^). Percentages of total lung D^b^-NP tetramer+ cells found in the parenchyma (top right panel) or circulation (bottom right panel) of each mouse. **D.** Representative flow plot (left panel) and quantitation (right panel) of D^b^-NP tetramer positive CD44^hi^ CD8^+^ memory T cells from the spleen. **E.** Representative flow plot of total lung CD8^+^ T cells showing D^b^-PA^+^ tetramer staining in the parenchymal (CD45 i.v.^-^) or circulation (CD45 i.v.^+^). Percentages of total lung D^b^-PA tetramer+ cells found in the parenchyma (top right panel) or circulation (bottom right panel) of each mouse. **F.** Representative flow plot (left panel) and quantitation (right panel) of D^b^-PA tetramer positive CD44^hi^ CD8^+^ memory T cells from the spleen. Data is representative of 1 of 2 repeats with 3 pairs each. * p<0.05 Student’s two-tailed t-test with unequal variance.

**Figure S4.**
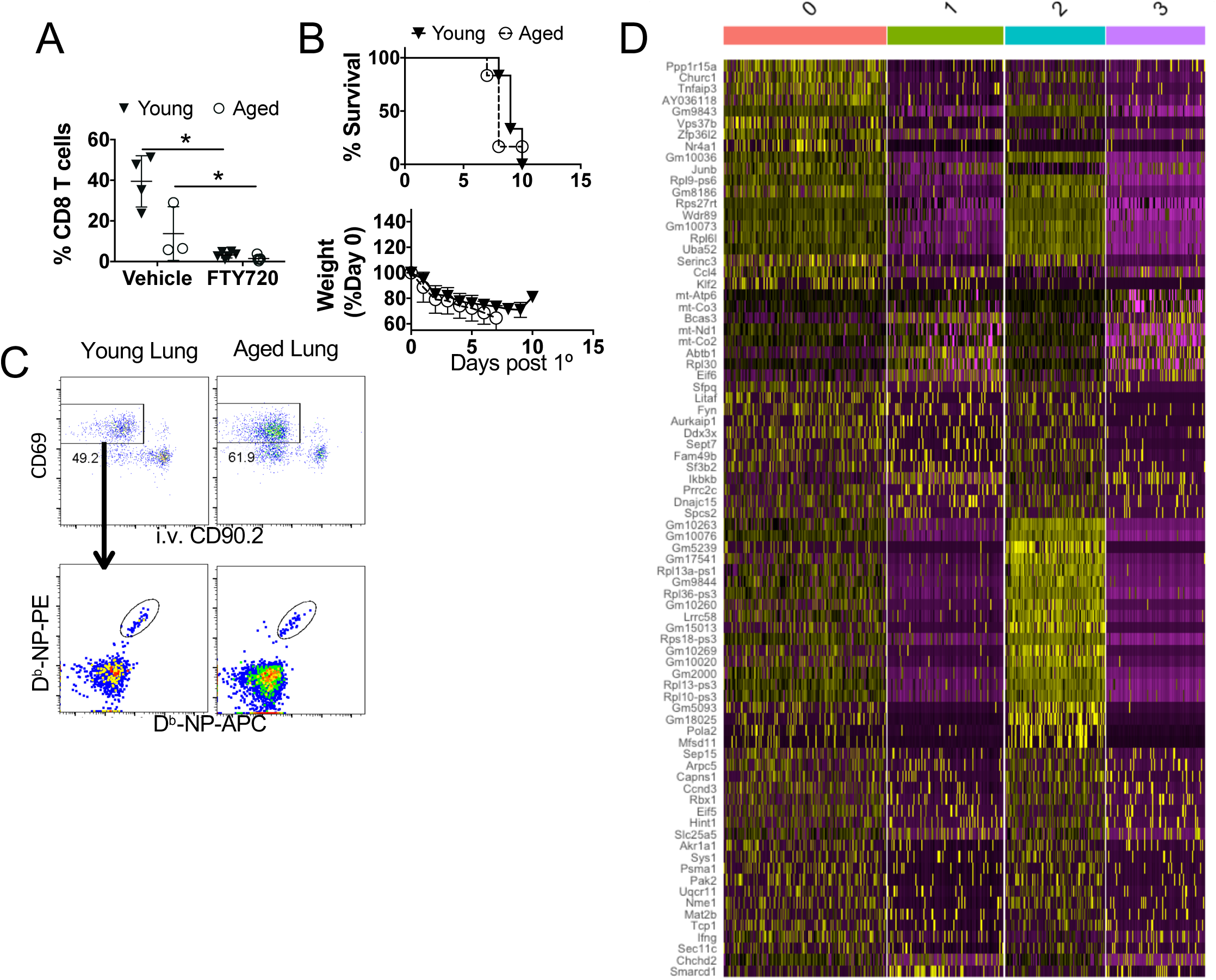
scRNA-seq on D^b^-NP T_RM_ cells from young or aged lungs. **A.** Percent of peripheral blood CD8 T cells in CD45^+^ white blood cells were evaluated 48 hours following FTY720 or vehicle treatment in young and aged memory mice. **B.** Survival (top) and weight loss (bottom) following lethal X31 infection of naive young or aged mice in the presence of FTY720. **C.** Sorting scheme for D^b^-NP specific T_RM_ cells. Following CD8 enrichment (Miltenyi kit), CD44^Hi^CD69^+^i.v.CD90.2^-^D^b^-NP-PE^+^/ D^b^-NP-APC^+^ cells were sorted from young (n=18) or aged (n=11) mice at 60 d.p.i. for scRNAseq. **D.** Heat map of top 20 DEGs by cluster for scRNA seq data. scRNAseq data is from a sorted pool of 18 young or 11 aged mouse lungs. **A & B** are each single replicates.

**Figure S5.**
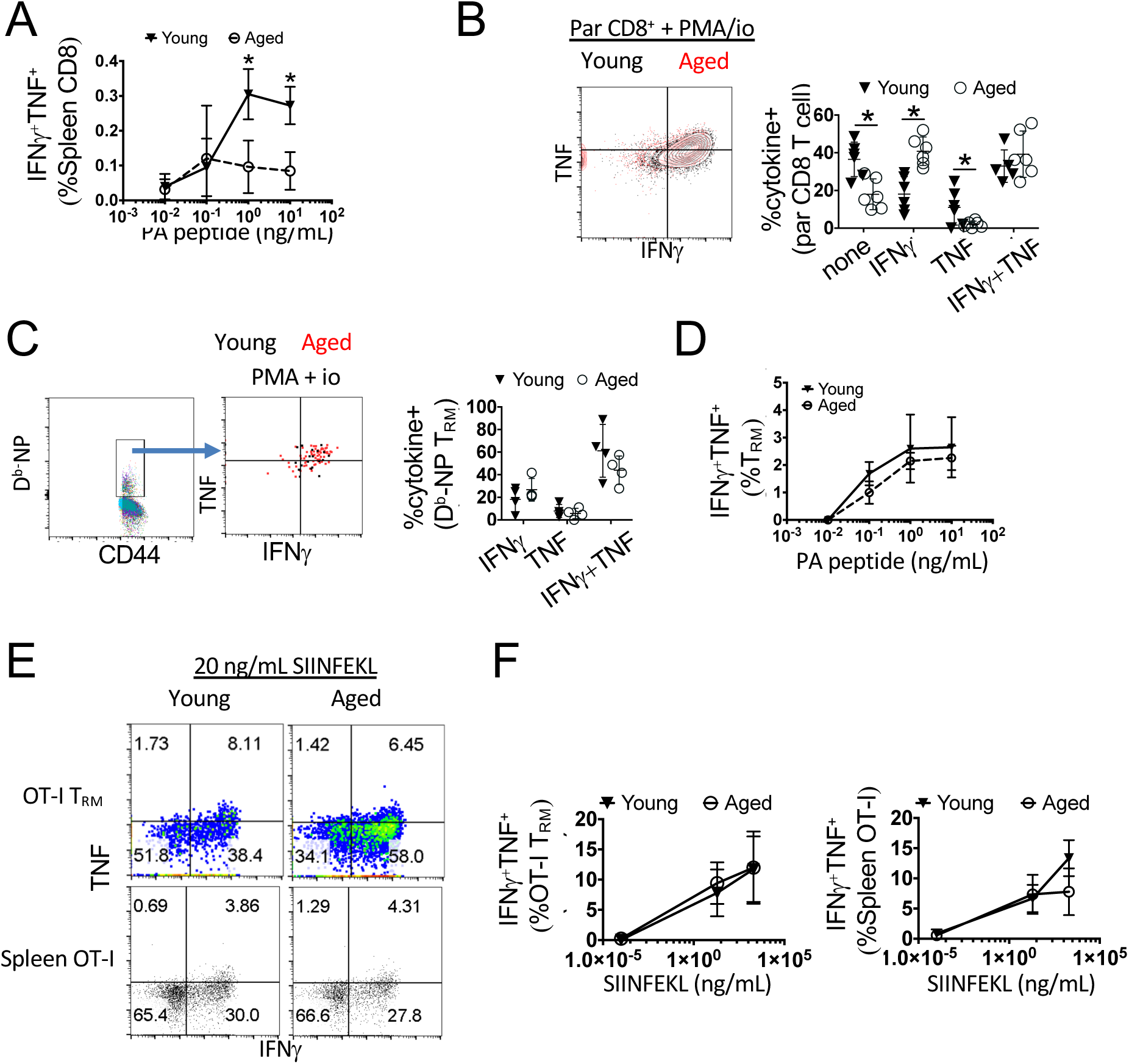
D^b^-PA or OT-I T_RM_ cells from aged hosts show equivalent of cytokine production as those of young hosts. Young or aged mice were infected with PR8. **A.** Percent of IFN-γ^+^ splenic CD8^+^ T cells following stimulation with increasing amount of NP 366-374 peptide (0.01 – 10 ng/mL) at 60 d.p.i. **B**. Representative plots (left) or percent of IFN-γ^+^ TNF^+^ lung-resident CD8^+^ T cells following stimulation with PMA/Ionomycin at 60 d.p.i. **C.** Representative plots (left) or percent of IFN-γ^+^ TNF^+^ lung-resident CD8^+^ D^b^-NP^+^ T cells following stimulation with PMA/Ionomycin at 60 d.p.i. **D.** Percent of IFN-γ^+^ TNF^+^ lung-resident CD8^+^ T cells following stimulation with increasing amount of PA 224-233 peptide (0.01 – 10 ng/mL) at 60 d.p.i. **E, F.** OT-I T cells (CD90.1^+^) from young donors were adoptively transferred into Young or Aged C57 BL6 mice 1 day prior to infection with PR8-OVA virus. Representative flow plots **(E)** or percent **(F)** of IFN-γ^+^ TNF^+^ lung-resident or splenic OT-I cells following stimulation with increasing amount of SIINFEKL peptide at 50 d.p.i. **A-D** are representative of 3 experiments each and **E & F** are representative of 2 experiments. * p<0.05 Student’s two-tailed t-test with unequal variance.

**Figure S6.**
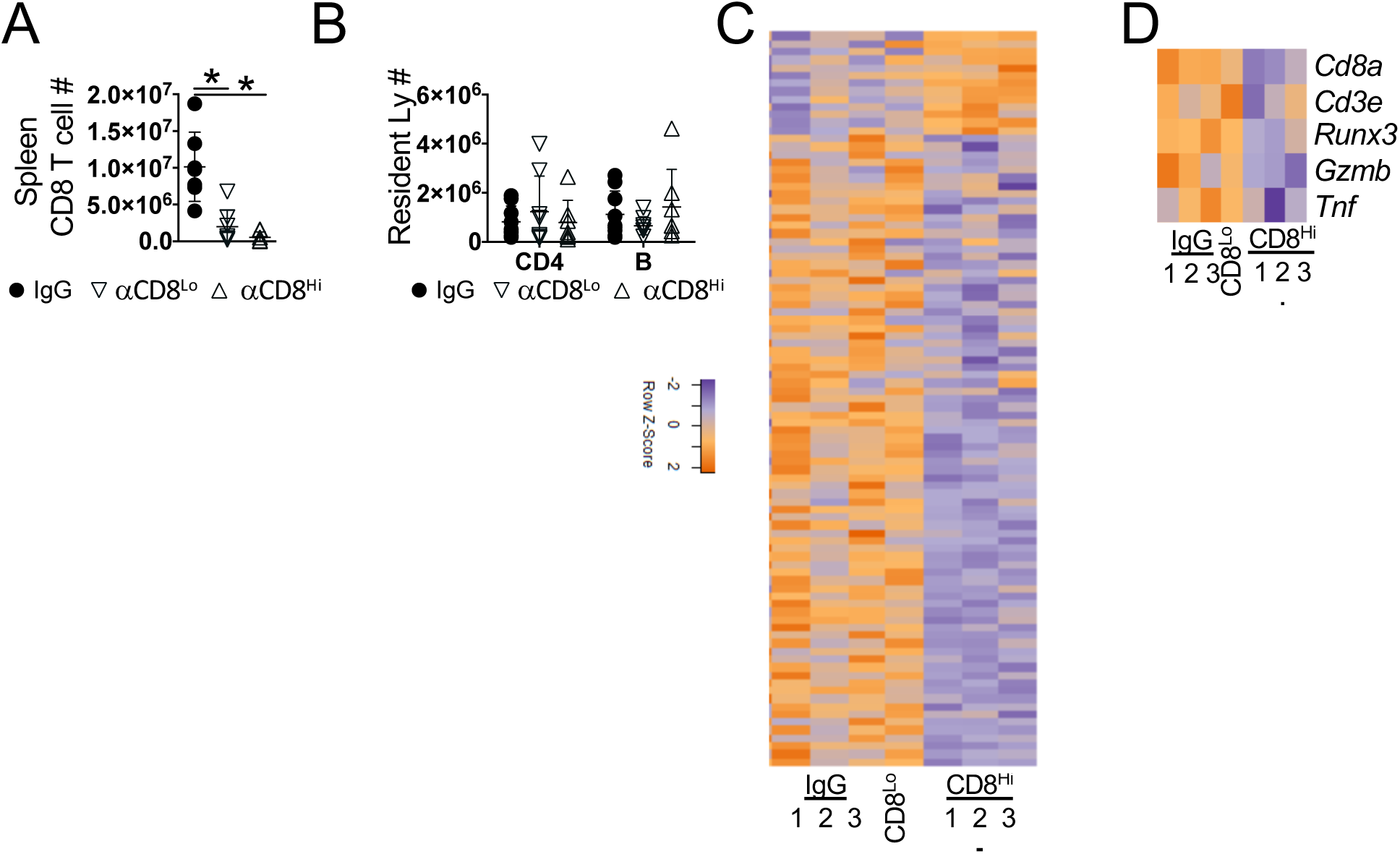
High dose of CD8 Ab treatment diminishes lung inflammatory responses. Aged or young C57BL/6 mice were infected with PR8. Mice received high or low dose of anti-CD8 treatment starting at day 21 as indicated. **A.** Splenic CD8^+^ T cell numbers in aged mice following CD8 Ab treatment. **B.** Resident CD4 and B cells were quantitated in the lungs. **C.** Complete set of DEGs (1.5 fold) from Nanostring data in Figure 6 G. **D.** Select nanostring DEGs showing CD8-related gene depletion after high, but not low dose Ab treatment in aged mice at 60 d.p.i. **A & B** are 2 pooled experiments, **C & D** are a single experiment in triplicate or pooled from 3 mice (αCD8^Lo^). * p<0.05 ANOVA with correction for multiple tests.

